# Fluid mechanics of mosaic ciliated tissues

**DOI:** 10.1101/2021.03.31.437829

**Authors:** Francesco Boselli, Jerome Jullien, Eric Lauga, Raymond E. Goldstein

## Abstract

In tissues as diverse as amphibian skin and the human airway, the cilia that propel fluid are grouped in sparsely distributed multiciliated cells (MCCs). We investigate fluid transport in this “mosaic” architecture, with emphasis on the trade-offs that may have been responsible for its evolutionary selection. Live imaging of MCCs in embryos of the frog *Xenopus laevis* shows that cilia bundles behave as active vortices that produce a flow field accurately represented by a local force applied to the fluid. A coarse-grained model that self-consistently couples bundles to the ambient flow reveals that hydrodynamic interactions between MCCs limit their rate of work so that when the system size is large compared to a single MCC, they best shear the tissue at low area coverage, a result that mirrors findings for other sparse distributions such as cell receptors and leaf stomata.

---

An indication of the importance of fluid mechanics in biology is the remarkable degree to which the structure of eukaryotic cilia has been conserved over the past billion years [1, 2]. These hairlike appendages provide motility to microorganisms [3, 4] but also direct fluid flow inside animals during development [5–7] and in mature physiology in areas from the reproductive system [8] to the brain [9]. The two extremes of this organisimal spectrum have a fundamental distinction. In unicellulars like *Paramecium*, cilia are uniformly and closely spaced on the cell surface [10], while in animals they are often grouped together in dense bundles on multiciliated cells (MCCs) [11] that are sparsely distributed on large epithelia, as in the trachea and kidney [12, 13]. This difference reflects the need in animal tissues to share surface area with cell types having other roles, such as mucus secretion.

The workings of cilia bundles and the significance of their sparse “mosaic” pattern for fluid transport have only begun to be investigated, primarily limited to *in vitro* or *ex vivo* studies [14–16]. Here we address the fluid mechanics of mosaic tissues using embryos of the amphibian *Xenopus laevis* in which, by analogy to human airways, cilia driven flow sweeps away mucus and trapped pathogens (Fig. 1). To date, the flow has served as a readout of cilia beating in the study of tissue patterning and cilia disorders [17–19]; here we take advantage of the geometry of *Xenopus* embryos to obtain side views of cilia bundles and quantify the flows they drive. As those cilia collectively sweep through cycles consisting of an extended “power” stroke and compact “recovery” stroke close to the surface [20], the flow within each bundle appears as an active vortex. While the flow driven by a single such vortex decays quickly with distance from the skin, a coarse-grained model shows that long range contributions of other bundles slows the decay of this *endogenous* flow and determines the shear stress at non ciliated cells. From measurements of beating changes induced by *exogeneous* flows, we determine linear response coefficients describing the coupling between forces applied by bundles and the flows they generate; we find that hydrodynamic interactions between MCCs lead to maximization at low area coverage of shear at the intervening tissue. These results thereby suggest an explanation for the low area coverages observed in nature.

**FIG. 1.**
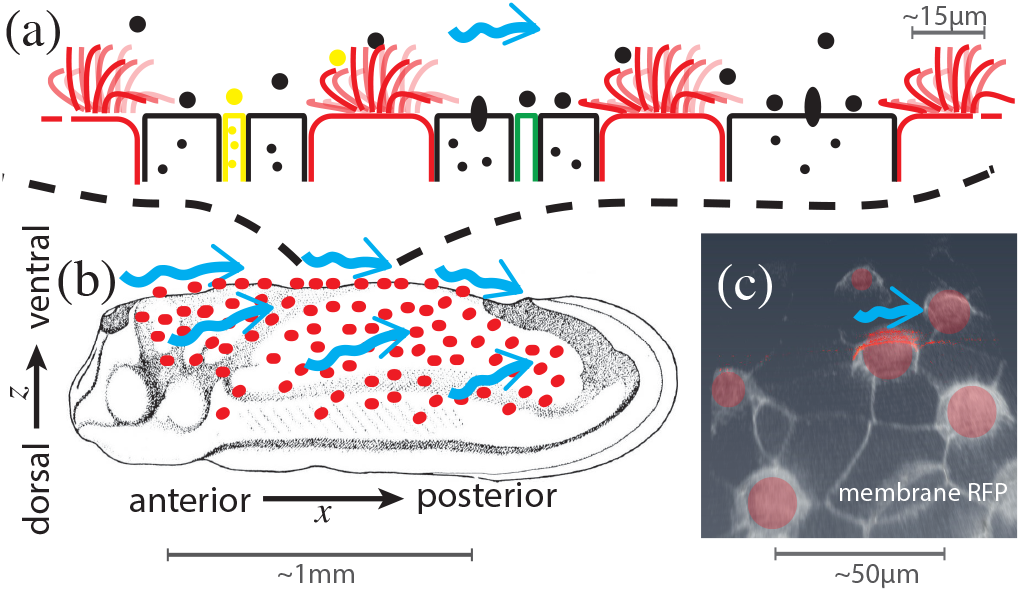
Ectoderm of embryonic *Xenopus laevis* at tailbud stages. (a) Schematic side view of MCCs (red) intermixed with secreting cells. (b) Location of MCCs across the embryo (adapted from [21, 22]) and cilia-driven flow (blue arrows). (c) Confocal image of cell membranes (stained by membrane-RFP), with MCCs segmented in red, in ventral region of skin.

The epidermis of *Xenopus* has strong similarities with human mucociliary epithelia. The sparsely located MCCs whence emanate hundreds of cilia that drive a homogeneous anterior-to-posterior, A-P or head-to-tail, flow (Fig. 1) are surrounded by non-ciliated cells secreting mucus-like material [23], including “goblet cells” that cover most of the tissue, mosaically scattered small cells [24, 25] secreting serotonin vesicles that modulate the cil-iary beat frequency [26], and ionocytes transporting ions important for homeostasis.

Wild-type *Xenopus laevis* embryos were obtained via *in vitro* fertilization [27, 28], and grown in 0 ×.1 Modified Barth’s Saline at room temperature (or 15° C to reduce the growth rate, if required). They were imaged at stage 28 [22] after treatment with a minimal dose of anaesthetic (∼ 0.01% Tricaine) to avoid twitching (without affecting cilia dynamics [19]). Embryos lie on one of their flat flanks at this stage, providing a side view of cilia bundles of ventral MCCs (Fig. 1), whose power strokes are in the A-P direction (left to right in figures) so cilia and the flows stay mostly within the focal plane.

In flow chamber experiments, embryos were perfused with a peristaltic pump while in a Warner Instruments chamber (RC-31A): a 4 mm × 37 mm channel cut into a 350 *µ*m thick silicon gasket sandwiched between two coverslips that keep the embryo in place by pressing against its sides. The dorsal part of the embryo was positioned closer to the chamber wall, the anterior region of interest was > 2 mm away, and the A-P axis parallel to the channel axis, the main direction of the perfusing flow. Brightfield images of cilia and 0.2 − 0.5 *µ*m tracers (mass fraction ~ 0.01%) were acquired at 2000 frames/s for ≥ 1 s by a high speed camera (Photron Fastcam SA3) on an inverted microscope (Zeiss Axio Observer) with a long distance 63× objective (Zeiss LD C-Apochromat). Images were filtered by subtracting their moving average. Flow fields **u** = (*u, v, w*) were estimated by Particle Image Velocimetry (PIVlab) and averaged over time.

We set the stage by summarizing the important length and time scales. MCCs are spaced apart by 40 − 80 *µ*m and uniformly distributed with average density P ≈2.6 × 10^−4^ *µ*m^−2^, giving an average spacing 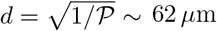. With 𝓁∼ 15 *µ*m 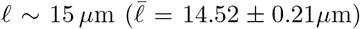 the cilia length, the average cellular area ∼ 287 ± 11 *µ*m^2^ is ∼ 𝓁^2^, which gives a coverage fraction *ϕ* = (𝓁*/d*)^2^ ∼ 0.07. The cilia on MCCs beat at a frequency *f* ∼ 20 − 30 Hz and during a power stroke their tips move a distance ∼ 2 𝓁 [Fig. 2(a)] in half a period, reaching speeds *V*_*c*_ ∼ 4*f𝓁* ∼ 1 mm/s, so the Reynolds number *ρV_c_ℓ*/*μ* (with *ρ* the density and *µ* the viscosity of water) is ∼ 0.01, well in the Stokesian regime. Using the fluid speed *u*_*v*_ ∼ 0.5 mm/s between vortices as typical of the periciliary region, the Péclet number *u*_*v*_ *𝓁/D* > 1 even for small molecules.

**FIG. 2.**
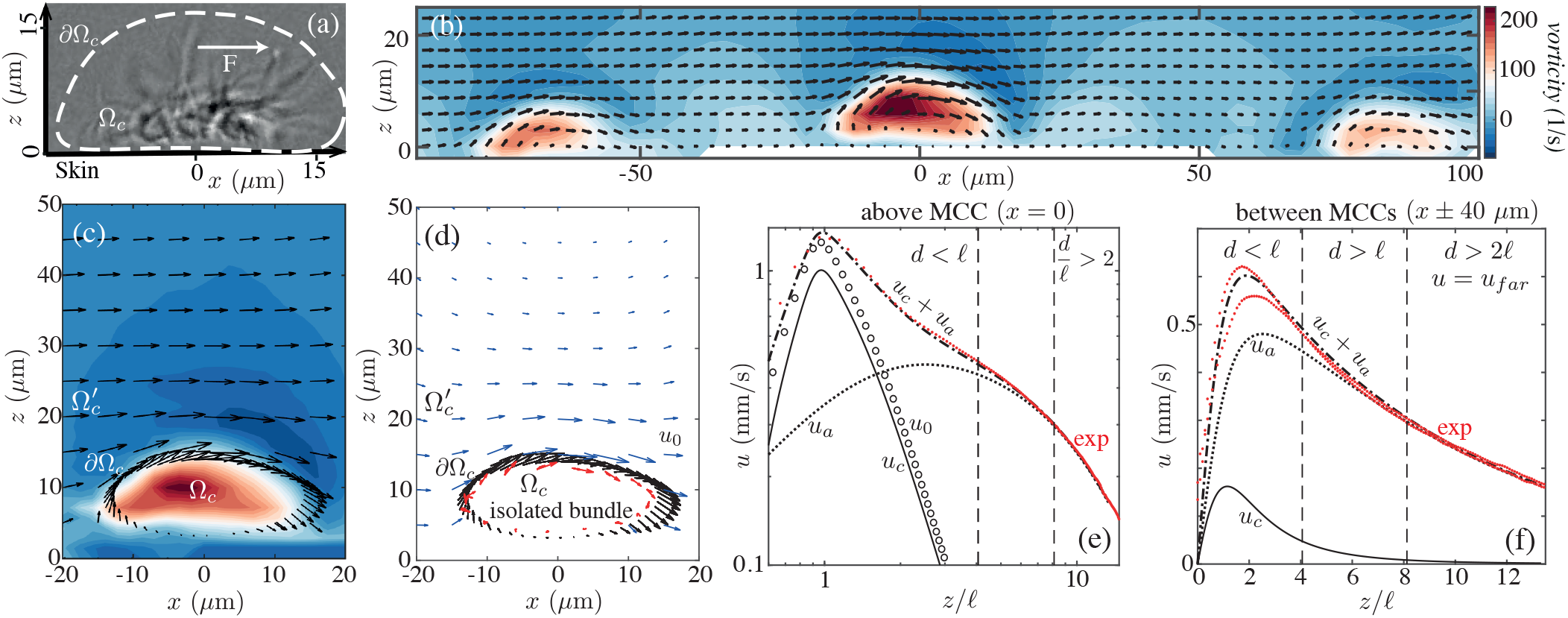
Flow field around multiciliated cells. (a) Lateral view of an MCC showing (dashed) path of cilia tips and applied force **F**. (b) Experimental velocity field and vorticity in a plane normal to skin near several MCCs, with *z* into fluid and *x* along flow. (c) Near an MCC, as in (b), with direction of cilia tip motion (black arrows) on *∂*Ω_*c*_. (d) Estimated flow field *u*_0_ for an isolated MCC (blue arrows): Stokeslets (red arrows) are used to fit velocity near cilia tips. Lateral velocity as a function of *z* at (e) *x, y* = (0, 0) and (f) (±40 *µ*m, 0) in experiment (exp) and theory, with *u*_0_ driven by an isolated bundle and *u*_*c*_ by a bundle exposed to the endogenous flow *u*_*a*_ (see also Figs. S1-S3 [28]).

Cilia within an MCC are not synchronized; their tips move in a tank-treading manner [28], generating vorticity ***ω*** ∥**e**_*y*_ perpendicular to the beating plane. Each MCC is thus an *active vortex*, as seen in Figs. 2(b,c). The vorticity can exceed ∼150 s^−1^ ∼2*V*_*c*_*/𝓁*, is colocalized with the cilia, and rapidly diffuses at larger *z* as the flow becomes parallel to the skin. Above non-ciliated cells between MCCs, there is a shear flow for *z* <*𝓁*, while further away (*z* ≳ *d*), the discreteness of the MCCs is washed out by viscosity and the horizontal velocity *u* is independent of *x* and falls off slowly with *z* [Figs. 2(e,f)].

The first step toward understanding the coupling between cilia beating and fluid flow involves quantifying the contribution of a single MCC. We introduce a boundary *∂*Ω_*c*_ enclosing the volume Ω_*c*_ of the active vortex [Fig. 2(a)], and extrude it in *y* ∈ [−10 *µ*m, 10 *µ*m], the measured size of the vortex. The Stokes equations in the complement 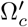, are solved using an envelope approach [29, 30] in which the dynamics of the cilia tips determine the flow, noting that the flow from an isolated cilium is well-approximated in the far field as that of a point force (Stokeslet) [31]. We position *N* Stokeslets at **s**_*n*_ = (*x*_*n*_, *y*_*n*_, *h*_*n*_) in Ω_*c*_ and find their strengths **f**_*n*_ = (*f*_*n,x*_, 0, *f*_*n,z*_) by fitting the velocity on *∂*Ω_*c*_ and a no-slip boundary at *z* = 0 [Fig. 2(d)]. This gives an estimate of the flow **u**_0_ driven by a bundle in an otherwise quiescent fluid. The *x*-components are of interest, as they alone contribute to net AP flow. For *z* > 2*l*, the component *u*_0_ falls off for *z ≫h* like the flow *u*_*s*_ (*z*) ∼3*Fh*^2^*/*4*πµz*^3^ above a single Stokeslet parallel to and a distance *h* above a no-slip wall [32] (Fig. S1 [28]). The data in Figs. 2(e,f) show a much weaker fall-off than *z*^−3^ for *z* > 2*l*. This slow decay arises from the long range contribution of more distant MCCs, as we now show.

The flow *u*_*s*_ due to a force *f*_*x*_ can be approximated by its far field limit. In cylindrical coordinates (*ρ, z*) centered at the bundle, 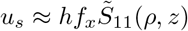, with

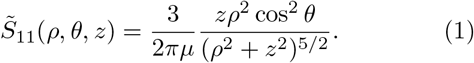

The Stokeslet contributions can be lumped into an effective force *F*_*c*_ = 𝓁^−1^ Σ_*n*_ *h*_*n*_*f*_*n,x*_, which, applied at *z* =𝓁, matches the far field 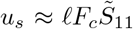. This also applies for an effective local moment (rotlet) Γ_*c*_ = 2 𝓁*F*_*c*_ [33].

The observed velocity *u*(*z*)**e**_*x*_ in the region *z* > *d* is independent of *x* (Fig. S2 [28]) and can be described most simply in a model of the skin as a uniform distribution of *x*-directed Stokeslets with area density 𝒫. Summing the contributions of all bundles up to a cutoff radius Λ that incorporates finite embryo size, we obtain

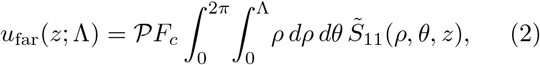

which has the scaling form *u*_far_ = *V G*(*z/*Λ), with

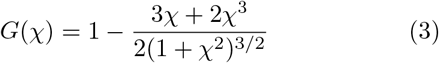

and *V* = 𝒫*F*_*c*_*l/µ. G* decreases monotonically from *G*(0) = 1 to *G*(∞) = 0. For any fixed *z*, as the organism size Λ → ∞, *χ* →0, giving a flow independent of *z* with speed *V* [34], while for any fixed lateral scale Λ, the asymptotic flow field vanishes as *z/*Λ → ∞. For the fitted parameters *V ≃* 0.64 mm/s and Λ = 300 *µ*m, *u*_*far*_ provides an almost perfect fit of the data in Figs. 2(e,f) for *z* > 2*d* (also Fig. S2 [28]). Direct summation of discrete Stokeslets on a lattice produces nearly identical results, validating the approximation of a continuous distribution for the far-field flow (Fig. S3 [28]). For this value of *V* and the observed density 𝒫, we have the farfield estimate *F*_*c*_ *≃* 159 pN. A slightly larger effective force *F*_0_ ≃ 200 pN is obtained by solely fitting the velocity at the cilia tips (Fig. 2d), which is to be expected as this approach does not account for the endogenous flow *u*_*a*_**e**_*x*_ to which a bundle is self-consistently exposed.

We estimate the endogeneous ambient flow *u*_*a*_ by sub-tracting from *u*_far_ the contribution from a single cilia bundle, taken as a distribution of radius *d/*2,

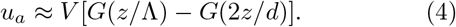

As *u*_*c*_ = *u*− *u*_*a*_ represents the contribution of a single bundle, we test the far-field estimate with a near-field fit of *u*_*c*_ in 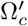 by means of Stokeslets within Ω_*c*_. Doing so for the volume (−15 < *x* < 15, −10 < *y* < 10, 0 < *z* < *d*), the superposition *u*_*c*_ + *u*_*a*_ gives an excellent fit to the data in Figs. 2(e,f), above and between bundles, with *F*_*c*_ *≃* 160 pN, nearly identical to the far-field estimate.

For comparison, the average lateral force generation over one cycle of beating (assuming only the power stroke contributes) can be estimated from resistive force theory [35] as *f* ∼ *ζ*_⊥_ 𝓁*V*_*c*_*/*12 ≃3 pN [28], where 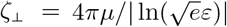 is the transverse drag coefficient for a slender filament of aspect ratio *ε* (for cilia, *ε* ∼ 75). We infer from the estimated *F*_*c*_ that the effective number of cilia contributing to the Stokeslet is ∼53, about half the typically ∼100 cilia in an MCC, reflecting force cancellations from phase shifts between cilia.

The fact that *F*_*c*_*/F*_0_ < 1 shows that it is necessary to incorporate the response of a cilia bundle to an ambient flow. We probed this experimentally by exposing the bundle to an exogenous shear flow 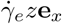 in a flow chamber (Fig. 3). When 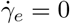, the cilia tips move with velocity *V*_0_ and drag the fluid, generating a negative shear rate 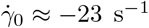. Pumping fluid in the same direction, the shear rate at the tips 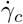 decreases linearly with the hydrodynamic load 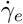. The corresponding velocity *V*_*c*_ tends to increase, but at a much slower rate. The rate of work above the cilia tip envelope 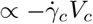 thus decreases almost at the same rate as 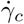, and for 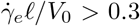 be-comes negative, consistent with a dissipative bundle.

**FIG. 3.**
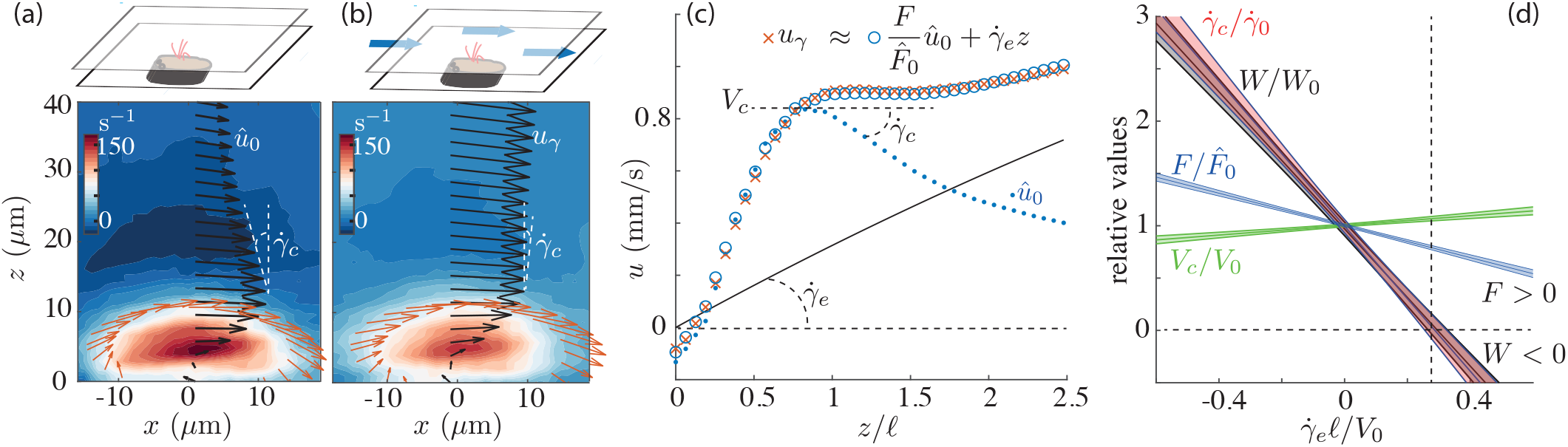
Response of a cilia bundle to shear flow. (a,b) Vorticity contours and velocity vectors before and during perfusion. (c) Velocities *û*_0_ and *u*_*γ*_, exogenous shear flow 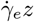, and linear combination 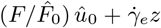 fitting *u*_*γ*_. (d) Linear fits of the variation of estimated force *F*, velocity *V*_*c*_ and shear rate 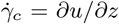 measured above cilia tips, and rate of work 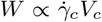 (overlaping 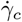), normalized by values at 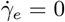. Shaded regions are 95% confidence intervals of averages over 10 samples.

The lateral velocity *u*_*γ*_**e**_*x*_ just above the bundle (*z* < 2.5𝓁) is well fitted by the linear combination *u*_*γ*_ ≈ *Cû*_0_ + *γ*_*e*_*z* (Fig. 3c) with 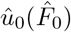 the profile at 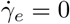. Note that 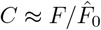, as confirmed for the above calculations where *u*_*c*_ ≈ (*F*_*c*_*/F*_0_)*u*_0_ for *z* >*𝓁* (Fig. S2(b) [28]). The slope of 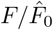 versus 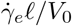 in Fig. 3, would be unity if the bundle dynamics were fully preserved (*V*_*c*_ = *V*_0_), and zero if the bundle’s force were constant. The measured slope 0.76 ± 0.06 confirms the resistive behavior and allows us to parameterize the coupling of the cilia to the ambient flow by the linear relation 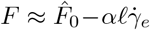, with 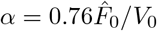.

To close the loop on a self-consistent coupling of the bundles to the flow, we now replace 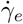 with the endogenous shear 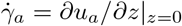, giving 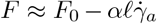, with *F*_0_ again the effective force applied by a bundle in an otherwise quiescent fluid. Since 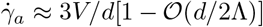 for large tissues, we have 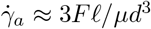, and thus

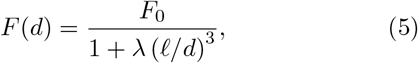

where the *λ* = 3*α/𝓁 µ* is the key parameter of the self-consistent theory. Using typical values 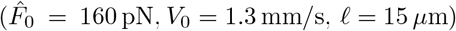, we find *λ* ≈ 18.6.

The relation (5) can be used to address several aspects of sparse cilia distributions [28]. The force applied to the wall per bundle is 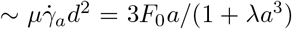, with *a* = *𝓁/d*, and has a maximum at *d*_*max*_ = (2*λ*)^1*/*3^ 𝓁 ≈ 50 *µ*m as does the contribution *FV* of a single bundle to the rate of work. The force 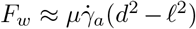 applied to the *non-ciliated* cells is maximal for *d* ≈54 *µ*m. Both values are in excellent agreement with those observed. Using the relation *ϕ* = (𝓁 */d*)^2^ we can express the wall force as a function of the coverage fraction *ϕ*,

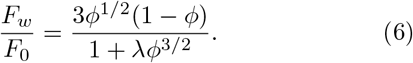

The contour plot of *F*_*w*_*/F*_0_ in the *ϕ* − *λ* parameter space in Fig. 4 shows that the optimum area fraction is a strongly decreasing function of *λ* and can reach values far below unity for *λ* ∼ 20, as in the present study. Extra endogenous loads 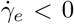, as expected for internal tissues, appear as an additional contribution to the ambient flow 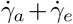. These loads will contribute to (5) as lower values of *λ*, and indeed, consistently larger, yet still low coverage fractions of the airways of several animals have been estimated to be 0.4 −0.5 [14], qualitatively consistent with Fig. 4. We infer that the observed mosaic patterns are close to optimal in terms of the clearing force applied to non-ciliated cells.

**FIG. 4.**
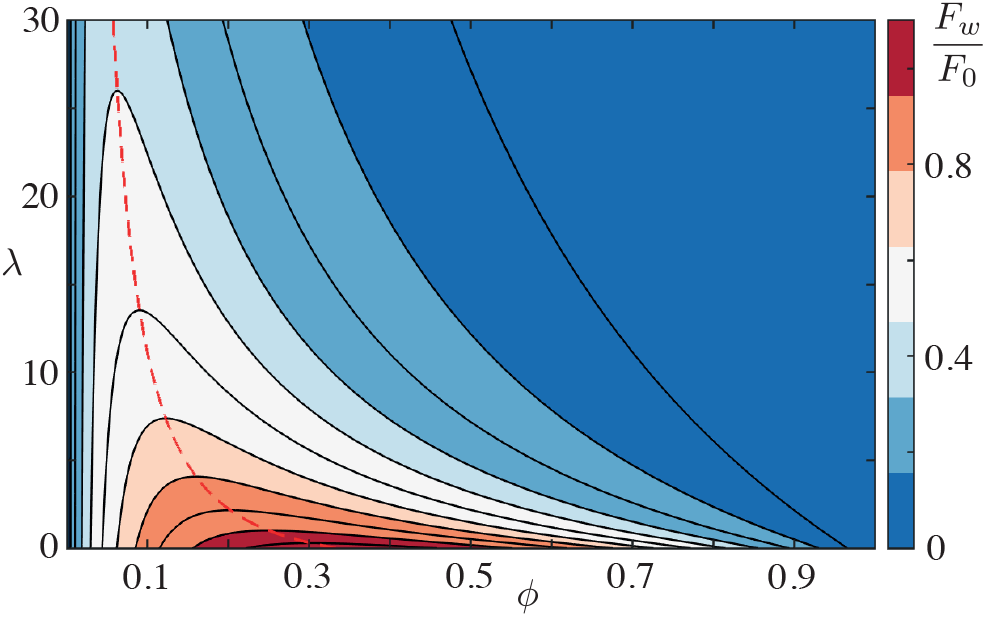
Force on non-ciliated cells in the self-consistent model. Contour plot of (6) in parameter space. Dashed line traces optimization ridge.

We close with comments on connections to other systems with sparse distributions of active elements. The force applied to the outer fluid by the cilia tips on the envelope *∂*Ω_*c*_ is equal and opposite to that applied to the skin, and we can simplify our results on the shearing of non-ciliated cells by reconsidering (2) as the flow of a patch of activity with given slip velocity *V* of radius Λ. The shear stress driving the flow is *τ*_Λ_ ≈3*µV/*2Λ, which we assume constant over a bundle. Setting Λ =𝓁 and integrating over a tissue with *N* bundles we obtain *J* ∼ *N 𝓁*^2^*τ*_*𝓁*_. By contrast, if we set Λ = *R*, the local shear stress is *τ*_*R*_ = 3*µV/*2*R* and the force over the entire surface is *J*_*R*_ ∼ *πR*^2^*τ*_*R*_. With *N* = *πR*^2^*ϕ/𝓁*^2^, the ratio

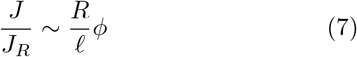

measures how well a distribution of non-interacting MCCs shears the surface relative to the collection. The linear scaling of (7) with *ϕ* is expected, but the large prefactor *R/𝓁* ∼ 20 (system size/MCC size) implies that *J/J*_*R*_ can approach unity for area fractions as low as *ϕ* ∼ 𝓁*/R* ∼ 5%. The form of this result mirrors one found by Jeffreys [36] for the evaporation rate from sparsely distributed leaf stomata, rediscovered years later [37] in the context of ligand binding to sparse cell receptors [38].

The results presented here suggest that long range hydrodynamic interactions between multiciliated cells allow efficient peri-ciliary transport at relatively low coverage, favoring the coexistence of multiple cell types in large tissues. This is likely just one aspect of more general mechanisms that maintain efficient transport in the up-scaling events marking the evolutionary transition from unicellular to larger multicellular systems.

This work was supported in part by Wellcome Trust Grant 101050/Z/13/Z) and Medical Research Council grant MR/P00479/1 (JJ), ERC Consolidator grant 682754 (EL), Wellcome Trust Investigator Award 207510/Z/17/Z, Established Career Fellowship EP/M017982/1 from the Engineering and Physical Sciences Research Council, and the Schlumberger Chair Fund (REG).

## SUPPLEMENTAL MATERIAL

This file contains additional experimental and calculational details.

### Embryo Culture

*Xenopus* embryos were prepared as described previously [S1]. Briefly, mature *Xenopus laevis* males and females were obtained from Nasco [S2]. Females were injected with 50 units of pregnant mare serum gonadotropin 3 days in advance and 500 units human chorionic gonadotropin 1 day in advance in the dorsal lymph sack to induce natural ovulation. Eggs were laid in a 1× MMR buffer (5 mM HEPES pH 7.8, 100 mM NaCl, 2 mM KCl, 1 mM MgSO_4_, 2 mM CaCl_2_, 0.1 mM EDTA). *Xenopus* embryos were cultured at room temperature or 15° C in the 0.1 × MMR until they reached stage 27*/*28. Experiments with embryos were performed at the late tailbud stages (stages 28-30, as describe in Faber and Nieuwkoop [S3]). Embryos were terminated humanely immediately following the experiments.

Our work with *Xenopus laevis* is covered under the Home Office Project License PPL 70/8591 and frog husbandry and all experiments were performed according to the relevant regulatory standard. All experimental procedures involving animals were carried out in accordance with the UK Animals (Scientific Procedures) Act 1986. Moreover, we only used surplus embryos for this study, to conform with the NC3Rs guidance to exploit the possibility to minimise the use of animals by sharing embryos with collaborators.

### Statistics

To fit any quantity *y* measured at a given hydrodynamic load *x*, we assume a linear relation *y* = *a* + *bx* and find 95% confidence intervals for the averages *ā* and 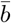 of the parameters *a*_*i*_ and *b*_*i*_ given by a least-square fit of the measured values (*x*_*i,m*_, *y*_*i,m*_) acquired for the *i*-th MCC. We have [S4]:

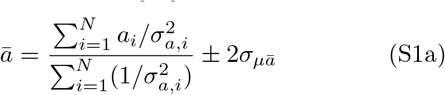

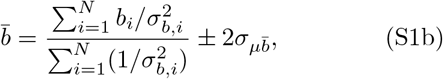

The standard errors for the mean parameters are the square roots of

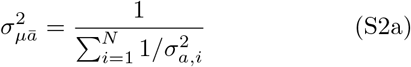

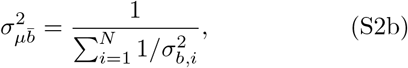

while the parameter variances for the *i*-th MCC are

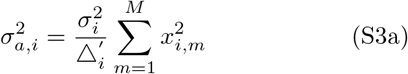

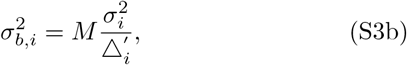

with

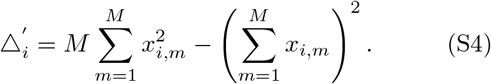

**FIG. S1.**
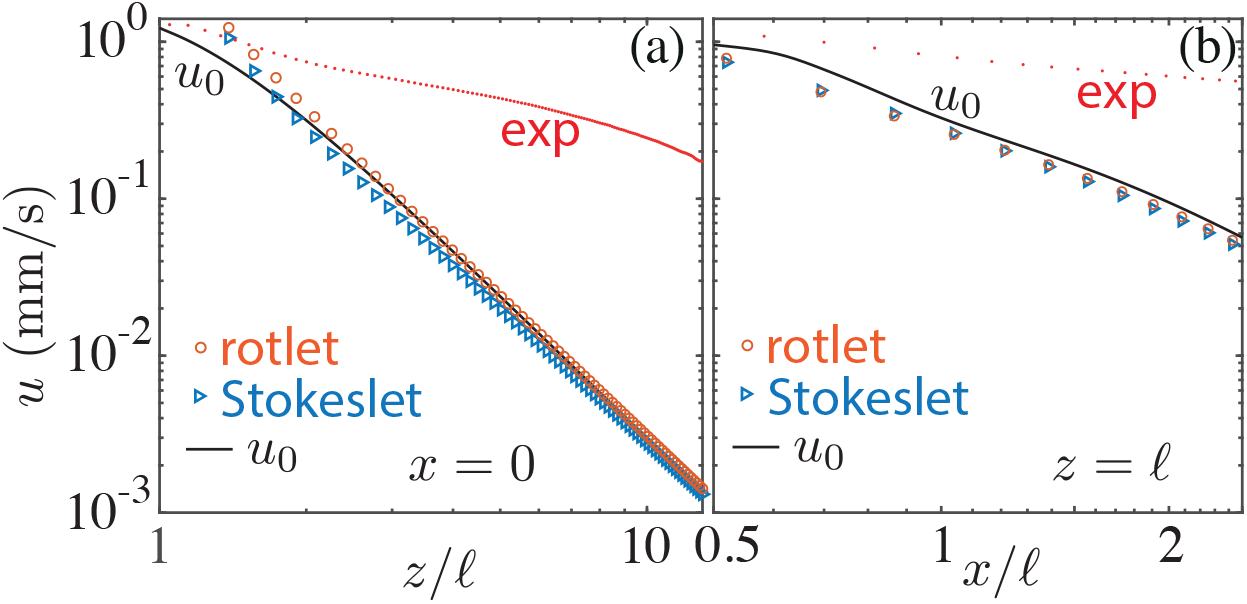
The contribution of a single bundle of cilia decays as a single effective force *F*_*c*_ or moment 2*𝓁F*_*c*_, while the measured flow profile decays much more slowly due to the contributions from other MCCs (exp). The lateral velocity *u*_0_, with reference to Fig. 2(d), is shown (a) above the bundle as a function of *z*, and (b) between bundles at *z* = *𝓁*, as a function of *x*.

**FIG. S2.**
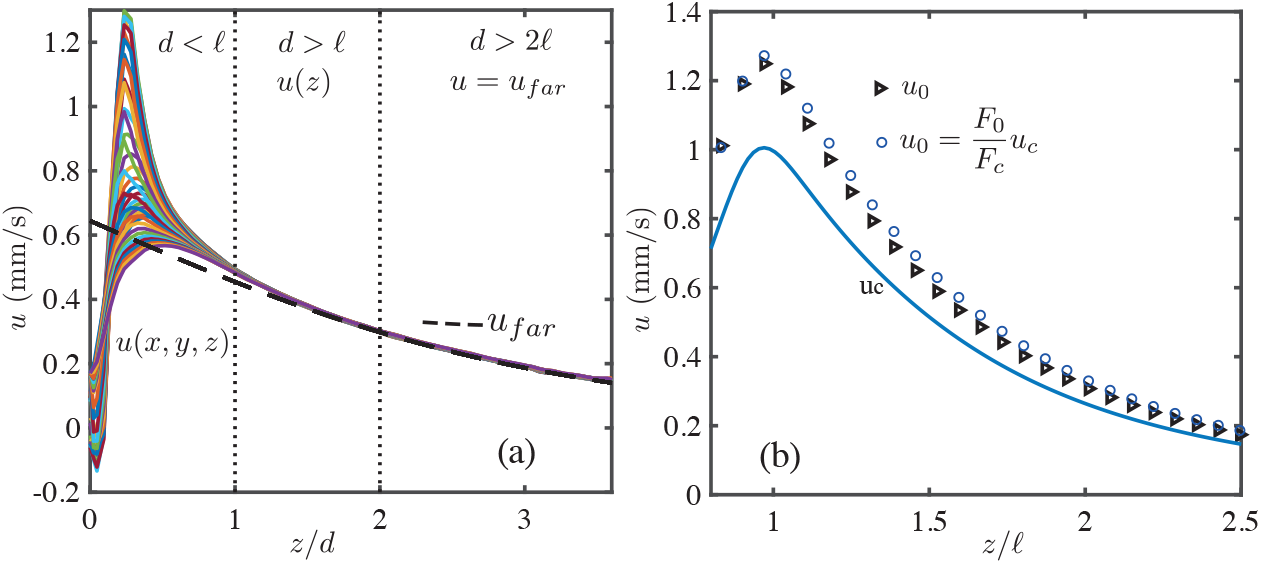
Supplement of Fig. 2. (a) Measured lateral velocity *u*(*z*) above and between bundles. For *z* > *d*, the *u*(*z*) becomes independent on *x*. For *d* > 2*d*, the *u*(*z*) matches the far-field model *u*_*far*_(*z*) of a uniform distribution of cilia. Different colours correspond to velocity profiles at several |*x*| < 35 *µ*m. (b) Linear dependance of the near-field *u*_*c*_ on the effective force *F*_*c*_, for *z* > *𝓁*.

The prime superscript indicates that the variance of the measurements 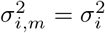 for the *i*-th MCC was assumed to be constant. It was estimated as the variance *s*^2^ of the sample population:

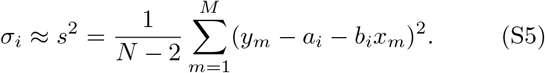

We normally imaged *M* = 4 conditions per MCC. For exceptions with *M* = 2, *s*^2^ could not be computed directly from (S5) and was assumed to equal the largest value from the other experiments.

### Fitting the near flow-field by the singularity method

The flow *u*_*c*_ driven by the cilia in Ω_*c*_ is modelled as the superposition of the flows arising from local point forces (Stokeslets) **f**_*n*_ applied at **s**_*n*_ ∈ Ω_*c*_:

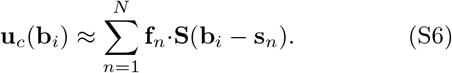

The tensor **S** is the well-known, exact solution for a Stokelset next to a no-slip plane at *z* = 0 [S5]. The values of **f**_*n*_ are found by fitting **u**_*c*_ at *M* collocation points **b**_*i*_, with *M* > 2*N* to avoid numerical instabilities [S6]. As no-slip boundary conditions **u**_*c*_ = 0 at *z* = 0 are implicitly satisfied, walls do not need to be discretized. The linear system (S6) is then simply recast in its matrix form **Af** = **u**_*b*_, with the 3*M* × 2*N* matrix

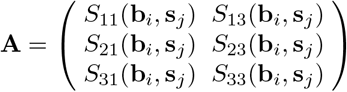

the 2*N* × 1 vector **f** = {*f*_1,1_, …, *f*_*N*,1_, *f*_1,3_, …, *f*_*N*,3_}, and the 3*M* 1 vector **u**_*b*_ = *u*_*c*,1_(**b**_*i*_), *u*_*c*,2_(**b**_*i*_), *u*_*c*,3_(**b**_*i*_). We then solve for **f** using the backslash operator of Matlab.

**FIG. S3.**
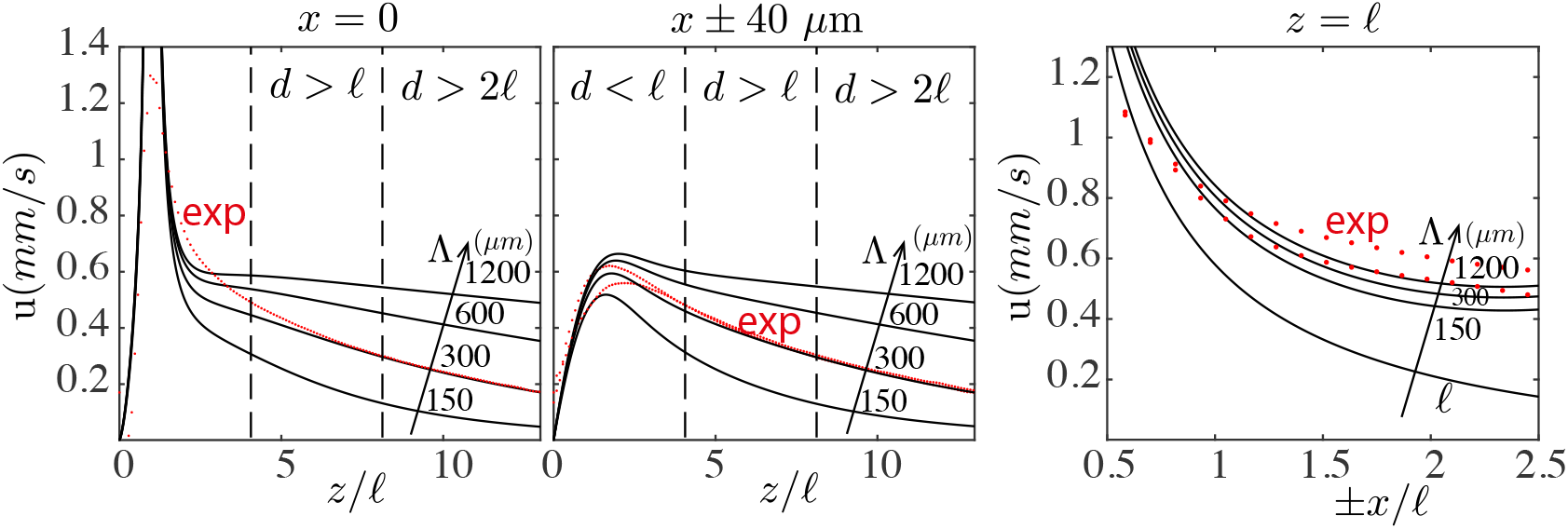
The lateral velocity of a two-dimensional array of Stokeslets of radius Λ ∼ 300, falls off as in experiments (exp), as shown for: (a,b) above and between bundles, respectively, as a function of *z*; (c) between bundles as a function of *x*. Results are obtained by direct summation of the exact solution of each Stokeslet, all of strength *F*_*c*_**e**_*x*_ and z-offset *𝓁*.

**FIG. S4.**
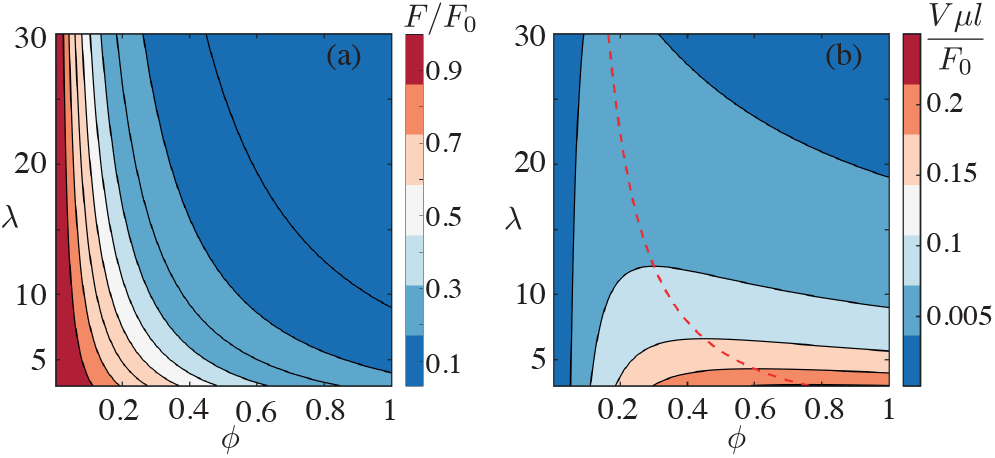
Supplement of Fig. 4. Contour plot of (a) the effective force *F* [Eq. (5)], and (b) the limit velocity *V*, in parameter space. Dashed red line in (b) traces ridge of optimization for *V*.

We set *f*_2_ = 0, assuming the solution to be symmetric in *y*. Once **f**_*n*_ are known, (S6) can be used to evaluate the fitted solution at any **x**. The flow field measured in the *y* plane is extruded by replications at 13 planes evenly spaced between − 10 *µ*m < *y* < 10 *µ*m. We used 15 Stokeslets for each plane about the fictitious boundary *∂*Ω_*c*_.

### Coarse-graining the bundle

The flow **u**_*c*_ driven by the Stokeslet in the bundle can be coarse-grained further, with a smaller number of Stokes flow singularities, moving away from the bundle. We compare the fitted flow **u**_*c*_, made up of *N* Stokeslets as discussed above, with the flow driven by the effective Stokeslet *F*_*c*_**e**_*x*_ applied at (0, 0, *𝓁*), and the effective rotlet 2 𝓁*F*_*c*_**e**_*y*_ applied at (0, 0, *𝓁/*2). They share the same far-field, reflecting the fact it is driven by an active vortex. The flow driven by the entire bundle decays as 1*/z*^3^, as for a single singularity, for *z* > 2 𝓁 (Fig. S1).

### Far Field fitting

The flow given by (2), is used to fit the PIV measurements for *z* > 2*d* (Fig. S2). The velocity *u*_*far*_(*z*; Λ) depends linearly on *V*, but not on *R*. For a given value of *R*, we find *V* by a linear least-squares fit of the data. We then simply repeat this linear fit for candidate values in the range 30 *µ*m < *R* < 1 mm, with increment Δ*R* = 10 *µ*m, and select the value of *R* that minimizes the *L*_2_ fitting error.

### Two-dimensional array of Stokeslets

Results similar to those presented in Fig. 2 for a uniform distribution of Stokelets can be obtained by positioning Stokeslets *F*_*c*_**e**_*x*_ on a lattice with cut-off radius Λ. Each element **s**_*ij*_ of the lattice is position at (*x*_*ij*_ = *id*_11_ + *jd*_12_, *y*_*ij*_ = *jd*_22_, *z*_*ij*_ = 𝓁). From confocal imaging of the closest neighboring cells of the bundle in Fig. 2, we estimate *d*_11_ ∼ 70 *µ*m, *d*_12_ = 40.5 *µ*m, *d*_22_ = 53*µm*.

Using the effective force *F*_*c*_ estimated by the near field fitting, and summing up the exact contribution of each MCC, we retrieve the slow decay rate observed *in vivo* [Figs. 2(e,f)] for Λ ∼ 300 *µ*m (Fig. S3). This is the same result found by fitting the far-field flow with Eq. (2).

### Resistive force theory estimate of the effective force applied by a single cilium

We adopt a simplified view of the power stroke of a cilium as a straight rod that pivots around its base. Let *s* ∈ [0, *𝓁*] be arclength along a cilium, with *s* = 0 at the base and *s* = 𝓁at the tip, and let *ϕ* be the angle between the cilium and the wall. The lateral component of the RFT force density at *s* is *f* ′ ∼ (*s/ 𝓁*)*ζ*_⊥_*V*_*c*_ sin *ϕ*, and the resulting far field velocity, given by Eq. (1), is proportional to *hf* ′*ds* with *h* = *s* sin *ϕ*. Accordingly, the effective force 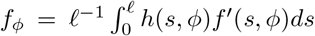 matches the overall far field 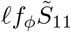 when applied at 𝓁. We obtain *f*_*ϕ*_ = sin^2^ *ϕ ζ*_⊥_ *𝓁V*_*c*_*/*3. Through the entire stroke, a cilium cycles through an angle Δ*ϕ* = 2*π*, and we assume that the recovery stroke does not contribute to the force, so *f*_*ϕ*_ = 0 for *π* < *θ* < 2*π*. Averaging *f*_*ϕ*_ gives the effective force 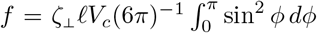 which gives the expression *f* = *ζ*_⊥_ 𝓁; *V*_*c*_*/*12 used in the main text.

### Additional results from the self-consistent model

The self-consistent model in Eq. (5) can be used to investigate several aspects of the phenomenology of cilia driven flow. The contour plots for the effective force *F* and the limit velocity *V* are shown in Fig. S4. *V* has a maximum at *d* = (*λ/*2)^1*/*3^𝓁≈ 32 *µ*m. The corresponding coverage fraction *ϕ* = (*λ/*2)^−2*/*3^∼ 0.22 is significantly larger than observed *in vivo*, confirming that the system is instead optimized for the wall force *F*_*w*_ (Fig. 4) discussed in the main text.

### Supplementary video

Movie of a cilia bundle and 0.2 *µ*m diameter tracers, acquired at 2, 000 frames/s, and shown at 30 frames/s. Some larger beads are also present to help visualize the flows.

## References

[1] G.J. Pazour, N. Agrin, J. Leszyk, and G.B. Witman, Pro-teomic analysis of a eukaryotic cilium, J. Cell Biol. 170, 103 (2005).

[2] C. Ainsworth, Tails of the unexpected, Nature 448, 638 (2007).

[3] D.R. Mitchell, The Evolution of Eukaryotic Cilia and Flagella as Motile and Sensory Organelles, In: Eukaryotic Membranes and Cytoskeleton. Advances in Experimental Medicine and Biology 607 (2007).

[4] R.E. Goldstein, Green algae as model organisms for biological fluid dynamics, Annu. Rev. Fluid Mech. 47, 343 (2015).

[5] T. Nakamura and H. Hamada, Left-right patterning: conserved and divergent mechanisms, Development 139, 3257 (2012).

[6] O. Thouvenin, L. Keiser, Y. Cantaut-Belarif, M. CarboTano, F. Verweij, N. Jurisch-Yaksi, P.L. Bardet, G. van Niel, F. Gallaire, Origin and role of the cerebrospinal fluid bidirectional flow in the central canal, eLife 9 (2020)

[7] R.R. Ferreira, A. Vilfan, F. Jülicher, W. Supatto, J. Vermot, Physical limits of flow sensing in the left-right organizer, eLife 6 (2017)

[8] L.J. Fauci and R. Dillon, Biofluidmechanics of reproduction, Annu. Rev. Fluid Mech. 38, 371 (2006).

[9] R. Faubel, C. Westendorf, E. Bodenschatz, and G. Eichele, Cilia-based flow network in the brain ventricles, Science 353, 176 (2016).

[10] H. Soares, B. Carmona, S. Nolasco, and L.V. Melo, Polarity in Ciliate Models: From Cilia to Cell Architecture, Front. Cell Dev. Biol. 7, 240 (2019).

[11] E.R. Brooks and J.B. Wallingford, Multiciliated cells: a review, Curr. Biol. 24, R973 (2014).

[12] Y. Liu, N. Pathak, A. Kramer-Zucker, and I.A. Drummond, Notch signalling controls the differentiation of transporting epithelia and multiciliated cells in the zebrafish pronephros, Development 134, 1111 (2007).

[13] A. Vasilyev, Y. Liu, S. Mudumana, S. Mangos, P.-Y. Lam, A. Majumdar, J. Zhao, K.-L. Poon, I. Kondrychyn, V. Korzh, and I.A. Drummond, Collective cell migration drives morphogenesis of the kidney nephron, PLOS Biology 7, e1000009.(2009).

[14] G.R. Ramirez-San Juan, A.J.T.M. Mathijssen, M. He, L. Jan, W. Marshall, and M. Prakash, Multi-scale spatial heterogeneity enhances particle clearance in airway ciliary arrays, Nat. Phys. 16, 958 (2020).

[15] M.-K. Khelloufi, E. Loiseau, M. Jaeger, N. Molinari, P. Chanez, D. Gras, and A. Viallat, Spatiotemporal organization of cilia drives multiscale mucus swirls in model human bronchial epithelium, Sci. Rep. 8, 2447 (2018).

[16] E. Loiseau, S. Gsell, A. Nommick, C. Jomard, D. Gras, P. Chanez, U. D’Ortona, L. Kodjabachian, J. Favier, and A. Viallat, Active mucus-cilia hydrodynamic coupling drives self-organization of human bronchail epithelium, Nat. Phys. 16, 1158 (2020).

[17] V.C. Twitty, Experimental studies on the ciliary action of amphibian embryos, J. Exp. Zool. 50, 319 (1928).

[18] G.A. Deblandre, D.A. Wettstein, N. Koyano-Nakagawa, and C. Kintner, A two-step mechanism generates the spacing pattern of the ciliated cells in the skin of Xenopus embryos, Development 126, 4715 (1999).

[19] M.E. Werner and B.J. Mitchell, Using Xenopus skin to study cilia development and function, Methods Enzymol. 525,191–217 (2013).

[20] C. Brennen and H. Winet, Fluid mechanics of propulsion by cilia and flagella, Ann. Rev. Fluid Mech. 9, 339 (1977).

[21] K. Karimi, J.D. Fortriede, VS Lotay, K.A. Burns, D.Z. Wang, et al., Xenbase: A genomic, epigenomic and transcriptomic model organism database, Nucleic Acids Research 46, (D1) (2018).

[22] J. Faber and P.D. Nieuwkoop, Normal Table of Xenopus laevis (Daudin) (Garland Publishing Inc., New York, 1994).

[23] S. Nagata, Isolation, characterization, and extraembryonic secretion of the Xenopus laevis embryonic epidermal lectin, XEEL, Glycobiology 15, 281 (2005).

[24] E. Dubaissi and N. Papalopulu, Embryonic frog epidermis: A model for the study of cell-cell interactions in the development of mucociliary disease, Disease Mod. Mech. 4, 179 (2011).

[25] E. Dubaissi, K. Rousseau, R. Lea, X. Soto, S. Nardeosingh, A. Schweickert, E. Amaya, D.J. Thornton, and N. Papalopulu, A secretory cell type develops alongside multiciliated cells, ionocytes and goblet cells, and provides a protective, anti-infective function in the frog embryonic mucociliary epidermis, Development 141, 1514 (2014).

[26] P. Walentek, S. Bogusch, T. Thumberger, P. Vick, E. Dubaissi, T. Beyer, M. Blum, and A. Schweickert, A novel serotonin-secreting cell type regulates ciliary motility in the mucociliary epidermis of Xenopus tadpoles, Development 141, 1526 (2014).

[27] E. Hörmanseder, A. Simeone, G.E. Allen, C.R. Bradshaw, M. Figlmüller, J. Gurdon, and J. Jullien, H3K4 methylation-dependent memory of somatic cell identity inhibits reprogramming and development of nuclear transfer embryos, Cell Stem Cell 21, 135 (2017).

[28] See Supplemental Material at http://link.aps.org/supplemental/xxxforfurtherdetailsandresults.

[29] J.R. Blake, Infinite models for ciliary propulsion, J. Fluid Mech. 49, 2 (1971).

[30] T.J. Pedley, D.R. Brumley, and R.E. Goldstein, Squirmers with swirl: a model for Volvox swimming, J. Fluid Mech. 798, 165 (2016).

[31] D.R. Brumley, K.Y. Wan, M. Polin, and R.E. Goldstein, Flagellar synchronization through direct hydrodynamic interactions, eLife 3, e02750 (2014).

[32] J.R. Blake, Note on the image system for a stokeslet in a no-slip boundary, Math. Proc. Camb. Phil. Soc. 70, 303 (1971).

[33] J.R. Blake and A.T. Chwang, Fundamental singularities of viscous flow. Part I: The image systems in the vicinity of a stationary no-slip boundary, J. Eng. Math. 8, 23 (1974).

[34] N. Osterman and A. Vilfan, Finding the ciliary beating pattern with optimal efficiency, Proc. Natl. Acad. Sci. USA 108, 15727 (2011).

[35] J. Gray and G.J. Hancock, The propulsion of sea-urchin spermatozoa, J. Exp. Biol. 32, 802 (1955).

[36] H. Jeffreys, XXX. Some problems of evaporation, Philos. Mag. 35, 270 (1918).

[37] H.C. Berg and E.M. Purcell, Physics of chemoreception, Biophys.J. 20, 193 (1977).

[38] R.E. Goldstein, Coffee stains, cell receptors, and time crystals: Lessons from the old literature, Physics Today 71, 32 (2018).

## References

[S1] E. Hörmanseder, A. Simeone, G.E. Allen, C.R. Brad-shaw, M. Figlmüller, J. Gurdon, and J. Jullien, H_3_K_4_methylation-dependent memory of somatic cell identity inhibits reprogramming and development of nuclear transfer embryos, Cell Stem Cell. 6, 135 (2017).

[S2] www.enasco.com.

[S3] J. Faber and P.D. Nieuwkoop, Normal Table of Xenopus laevis (Daudin) (Garland Publishing Inc., New York, 1994).

[S4] P.R. Bevington, D.K. Robinson, Data Reduction and Error Analysis for the Physical Sciences (McGraw-Hill, 1993), 3rd edition.

[S5] J.R. Blake, Note on the image system for a stokeslet in a no-slip boundary, Math. Proc. Camb. Phil. Soc. 70, 303 (1971).

[S6] F. Boselli, D. Obrist, L. Kleiser. A multilayer method of fundamental solutions for Stokes flow problems, J. Comput. Phys. 231, 18 (2012).

